# NanoSplicer: Accurate identification of splice junctions using Oxford Nanopore sequencing

**DOI:** 10.1101/2021.10.23.465402

**Authors:** Yupei You, Michael B. Clark, Heejung Shim

## Abstract

**Motivation:** Long read sequencing methods have considerable advantages for characterising RNA isoforms. Oxford nanopore sequencing records changes in electrical current when nucleic acid traverses through a pore. However, basecalling of this raw signal (known as a squiggle) is error prone, making it challenging to accurately identify splice junctions. Existing strategies include utilising matched short-read data and/or annotated splice junctions to correct nanopore reads but add expense or limit junctions to known (incomplete) annotations. Therefore, a method that could accurately identify splice junctions solely from nanopore data would have numerous advantages.

**Results:** We developed “NanoSplicer” to identify splice junctions using raw nanopore signal (squiggles). For each splice junction the observed squiggle is compared to candidate squiggles representing potential junctions to identify the correct candidate. Measuring squiggle similarity enables us to compute the probability of each candidate junction and find the most likely one. We tested our method using 1. synthetic mRNAs with known splice junctions 2. biological mRNAs from a lung-cancer cell-line. The results from both datasets demonstrate NanoSplicer improves splice junction identification, especially when the basecalling error rate near the splice junction is elevated. Our method is implemented in the software package NanoSplicer, available at https://github.com/shimlab/NanoSplicer.

## 1 Introduction

Splicing is an essential mechanism in eukaryotic cells that removes introns from pre-mRNAs to create mRNA. Alternative splicing varies which sequences are intronic and exonic, enabling a single gene to produce multiple mRNA products (isoforms). Almost 95% of human genes (Pan *et al*., 2008) and 60% of Drosophila genes (Graveley *et al*., 2011) undergo alternative splicing, creating a diverse set of transcript isoforms whose expression can control cell functions in a particular condition or developmental stage. Short-read sequencing technologies (e.g., Illumina sequencing) have been successfully used to identify and quantify local splicing events, such as exon skipping. However, their read lengths (~ 150 nt) are much shorter than typical transcript lengths, making it difficult to combine each splicing event and identify the fulllength isoform(s) present (LeGault and Dewey, 2013; Steijger *et al*., 2013; Tang *et al*., 2020; Alqassem *et al*., 2021; Parker *et al*., 2021). As such our understanding of the isoform repertoire expressed in different organisms and those that control cell functions remains incomplete.

Nanopore sequencing by Oxford Nanopore Technologies (ONT) is a long-read sequencing method that can connect splicing events by sequencing full-length transcripts (Bolisetty *et al*., 2015; Byrne *et al*., 2017). Nanopore sequencing works by recording changes in electrical current when a DNA or RNA molecule traverses through a pore. This raw signal, (known as a *squiggle*) is then basecalled by computational methods, yielding reads that can cover the entire transcript and identify the expressed isoform. However, nanopore reads have a considerably higher basecalling error rate (~5-10%) and lower throughput compared to short reads, making their analysis challenging. In particular, the former makes read mapping near splice sites difficult (Tang *et al*. (2020); Volden *et al*. (2018); Weirather *et al*. (2017)) making it challenging to distinguish real splice junctions from mapping errors (Fig. 1A). Incorrect detection of splice junctions results in the identification of non-existent isoforms and omission of real isoforms, which inhibits the study of encoded proteins and isoform functions. In this paper, we develop a method to accurately identify splice junctions using nanopore sequencing, the performance of which is independent of sequencing throughput.

**Fig. 1.**
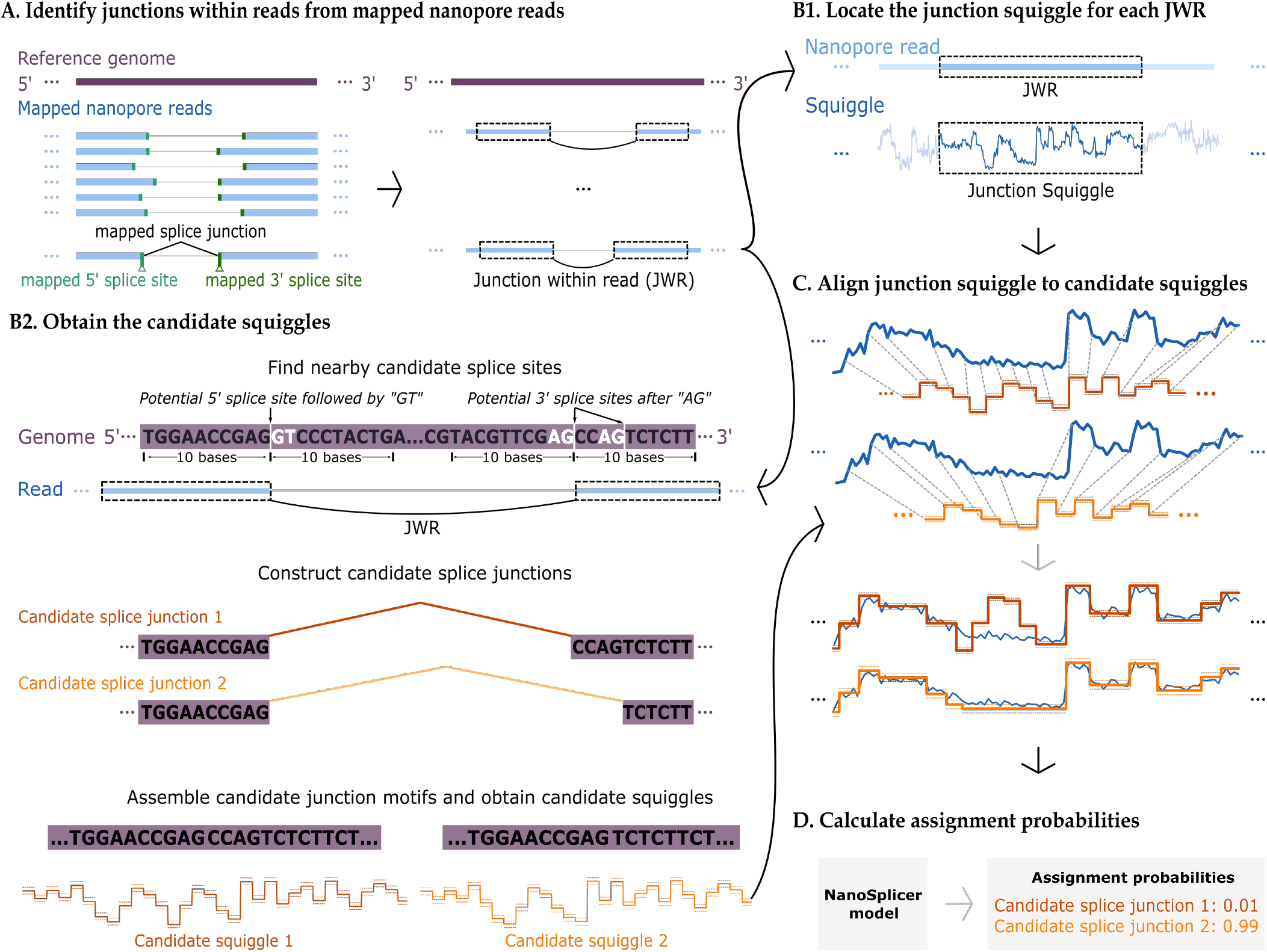
NanoSplicer workflow: A. Identify junctions within reads (JWRs). The left panel shows an example of inconsistently mapped splice junctions in nanopore reads, which may require correction. A splice junction refers to a pair of 5’ and 3 splice sites, which are the boundaries between introns and exons (shown in green in the figure). Right panel: NanoSplicer locates JWRs in mapped nanopore reads. The two dotted boxes connected by a black curve show a JWR, which is a subsequence of the read that is split and mapped to different exons. B1: Identification of *junction squiggles*. A basecalled nanopore read and its matched raw squiggle are aligned and the portion of the squiggle corresponding to the JWR (dotted boxes) is obtained. B2: Prediction of candidate squiggles. NanoSplicer identifies all possible canonical (“GT-AG”) splice junctions within 10 bases of the mapped splice sites. Possible 5’ and 3’ splice site nucleotides shown in white. Two candidate splice junctions (1 and 2) are shown (red and orange lines). Candidate junction motifs surrounding the splice junctions are then obtained using the reference genome and candidate squiggles for these motifs predicted with Tombo (Stoiber *et al*., 2016). Candidate squiggles include predicted mean current (solid line) +/- 1 standard deviation (dotted line). C: Alignment of candidate and junction squiggles. Top: The junction squiggle (blue) is aligned to each candidate squiggle (red/orange) using dynamic time warping. Dotted lines show which locations of the two squiggles are aligned. Bottom: Each current measurement in the junction squiggle (blue) is shown vertically aligned with its corresponding mean-standard deviation in the candidate squiggle. D: The NanoSplicer model provides assignment probabilities for each candidate by quantifying the squiggle similarity of each alignment.

Several authors have developed methods that correct splice junctions from mapped long reads (e.g., Wyman and Mortazavi (2019); Kovaka *et al*. (2019); Kuo *et al*. (2020); Tang *et al*. (2020); Parker *et al*. (2021)). Methods such as FLAIR (Tang *et al*., 2020) and TranscriptClean (Wyman and Mortazavi, 2019) require a set of splice junctions, either from annotations or from matched short reads, to be provided to guide their corrections. However, suitable annotations may not be available, for example, in a non-model organism or in a disease causing altered mRNA processing, while matched short-read sequencing increases costs. Furthermore, in some circumstances, short-reads covering whole transcripts cannot be generated. For example, nanopore sequencing is now being performed on single cells using the popular 10x Genomics platform, however, matched 10x short reads only cover transcript 3’ ends (Lebrigand *et al*., 2020). Other methods such as StringTie2 (Kovaka *et al*., 2019), TAMA (Kuo *et al*., 2020) and 2passtools (Parker *et al*., 2021) use information from other reads (e.g., nearby splice junctions supported by high read counts) to guide splice junction correction. A limitation with this approach is that it can lead to the replacement of rarer splice junctions with those from more highly expressed isoforms, causing less abundant junctions to go undetected (see Supplementary section 2.13 for an example). Moreover, methods that depend on other reads are not well suited to relatively low throughput nanopore datasets, where many isoforms, particularly from lowly expressed genes, have few reads.

The poor performance of nanopore read mapping near splice sites is largely due to basecalling errors, which arise when basecalling methods misinterpret the raw signal squiggles. Motivated by this, here we propose a method, NanoSplicer, that exploits the information in the squiggles to improve splice junction identification. The key idea is to identify, for each splice junction, which of the squiggles predicted from potential splice junction sequences best matches the observed junction squiggle. This “squiggle matching” idea has been successfully applied to map raw signals to a reference genome (Loose *et al*., 2016; Kovaka *et al*., 2021; Zhang *et al*., 2021), and we adapt this idea to develop a method for splice junction identification. By using the squiggle corresponding to each read, NanoSplicer does not require annotations or matched short reads and its performance is not affected by other reads and is independent of read depth, enabling it to identify rare splice junctions. We demonstrate the improved performance of NanoSplicer compared to the initial mapping using both synthetic and real data. Our method is implemented in the software package NanoSplicer, available at https://github.com/shimlab/NanoSplicer.

## 2 Methods

We developed NanoSplicer to accurately identify splice junctions using nanopore sequencing data. It takes as input mapped nanopore reads, their squiggles and a reference genome sequence. For each read it outputs lists of candidate splice junctions and the assignment probabilities quantifying the support for each of the candidates. Fig. 1 shows an overview of the NanoSplicer workflow. It consists of the following steps.

a. Locate subsequences in the mapped reads which split and map to different exons, supporting potential splice junctions. We refer to these subsequences as *junction within reads* (*JWRs*). See Supplementary section 1.1 for further details regarding JWR identification. For each JWR, we improve splice junction identification as follows:

Obtain the section of the squiggle corresponding to the JWR location, referred to as a *junction squiggle*.
Construct a list of candidate splice junctions, and predict an expected squiggle for each candidate, referred to as a *candidate squiggle*.
b. Align the junction squiggle to each of the candidate squiggles.
c. Use the NanoSplicer model to quantify the support for each candidate squiggle (assignment probability).

NanoSplicer also allows users to provide additional information to guide the choice of candidate splice junctions in step B2 (see section 2.2).

We discuss steps B-D in detail in the following sections.

### 2.1 Obtaining a junction squiggle

For each JWR, we obtain its *junction squiggle*, i.e., the squiggle section corresponding to the location of the JWR, as follows. First we use the “resquiggle” tool in Tombo (Stoiber *et al*., 2016) to align the nanopore read containing the JWR with its squiggle. Tombo performs the alignment by assigning current measurements in the squiggle to each base of the read. Then, we extract the part of the squiggle aligned to the JWR.

Tombo normalises squiggles during the alignment to remove systemic differences in shift (median value) and scale between squiggles (https://nanoporetech.github.io/tombo/resquiggle.html#signal-normalization). This normalisation enables the resulting junction squiggles to be comparable to candidate squiggles in sections 2.3 and 2.4. See Supplementary section 1.3 for additional squiggle preprocessing.

Basecalling errors create challenges in aligning current measurements to bases within reads and therefore in identifying the squiggle region corresponding to the JWR. However, matching over longer regions allows sub-regions with good alignment to be identified and the approximate position of the JWR to be specified. To implement this we take ~50 nt of the read as the JWR region, which allows us to identify the corresponding squiggle region even if exact base-current alignment for each nucleotide is not obtained. Rare cases where this process still identifies an incorrect region of the squiggle are filtered out (see section 2.4.4).

### 2.2 Obtaining candidate squiggles

We obtain candidate squiggles by first constructing a list of candidate splice junctions, and then for each candidate, identifying a candidate *junction motif* and predicting its expected *candidate squiggle* (Fig 1B2). We discuss each step in this section; see Supplementary section 1.2 for further details.

#### Candidate splice junctions

NanoSplicer provides multiple options to facilitate the selection of candidate splice junctions for each JWR, including a variety of inputs. This allows users to incorporate pre-existing information regarding splice junction usage (if available). By default NanoSplicer will select:

1. The splice junction supported by the JWR (mapped splice junction).
2. Nearby canonical splice junctions. We define these as introns that start with GT and end with AG (Fig. 1B2), a motif present in over ~99%. of mammalian splice junctions (Burset *et al*., 2000).

Inputs for each JWR can also include:

1. Annotated splice junctions.
2. User defined list of candidate splice junctions (e.g., from short-read sequencing).
3. Nearby splice junctions supported by other mapped reads (above a user-specified read count threshold).

Unless stated otherwise, we used the default option to choose candidate splice junctions in this paper. This allows NanoSplicer to identify splice junctions solely from the long-read data and does not require prior annotations or information from other reads. For ‘nearby canonical splice junctions’, NanoSplicer identified all GT and AG sequences within 10nt of the mapped 5’ and 3’ splice sites respectively and included the splice junctions these would create as candidates.

#### Candidate junction motifs

Once we construct a list of candidate splice junctions, we assemble a *junction motif* for each candidate by connecting sequences from each side of the candidate splice junction using the reference genome. Each candidate junction motif for a JWR extends 5’ and 3’ from the candidate splice junction to a common location. This ensures each candidate has the same nucleotide sequence (and squiggle signal) at the beginning and end, ensuring differences between candidate squiggles are solely due to the various splice junctions utilised.

#### Candidate squiggles

We predict a *candidate squiggle* for each candidate junction motif using an “expected current level model” in Tombo (Stoiber *et al*., 2016). This model provides the mean and standard deviation of the current level for each nucleotide in a candidate junction motif (https://nanoporetech.github.io/tombo/model_training.html describes how Tombo computes these means and standard deviations). The *candidate squiggle* can then be visualised by fitting a line through the sequence of means for each nucleotide.

### 2.3 Aligning the junction squiggle to each candidate squiggle

For each JWR, we now have its junction squiggle (section 2.1) and candidate squiggles (section 2.2). Before measuring the similarity between each candidate and junction squiggle, we first align them, i.e., assign current measurements in the junction squiggle to each mean and standard deviation in the candidate squiggle, so that their time axes are comparable (Fig 1C). We adapt Dynamic Time Warping (DTW) (Sakoe and Chiba, 1978) to align the two squiggles. DTW is an efficient algorithm for aligning two sequences which may vary in speed; see Keogh and Ratanamahatana (2005) for background on DTW. Supplementary section 1.4 describes our implementation of DTW which makes the following modifications.

1. We treat the junction squiggle as observations from a model that has the means and standard deviations of each candidate squiggle as parameters. Then, we use the support in the junction squiggle for the model as a measure of similarity in DTW.
2. A single observation has only one mean-standard deviation in a model. Thus, we assign each measurement in the junction squiggle to only one mean-standard deviation in the candidate squiggle.
3. In practice, the start and end of the junction squiggle may not perfectly match that of the candidate squiggles. Thus, we include more nucleotides at each side of the candidate junction motif as a buffer, and then allow the junction squiggle to be aligned to a part of the candidate squiggle.
4. The junction squiggle alignment is expected to cover most nucleotides in the candidate junction motif. Thus, we prevent current measurements in the junction squiggle from being aligned to only a small proportion of the candidate squiggle.

### 2.4 NanoSplicer model: accurate identification of splice junctions

Suppose, for a given JWR, we have its junction squiggle, *M* candidate squiggles, and *M* alignments, each of which aligns the junction squiggle to each candidate squiggle. Let ***x*** = (*x*_1_,…, *x_K_*) denote a junction squiggle with length *K*, where *x_k_* is the *k*-th current measurement. The *m*-th candidate squiggle, ***c***^*m*^, is the sequence of the mean-standard deviation of the current level for each nucleotide in its junction motif. The *M* alignments can be represented by an *M* × *K* matrix ***A*** = [*a_mk_*], where *a_mk_* indicates the index of the mean-standard deviation in ***c***^*m*^ where *x_k_* is aligned. Fig. S5 provides a toy example.

#### 2.4.1 Junction squiggle segmentation

Motivated by basecalling methods (Rang *et al*., 2018), we partition ***x*** into multiple segments, combine noisy measurements of ***x*** into a more stable summary value (e.g., mean, median) at each segment, and use the summary values as data in our NanoSplicer model (section 2.4.2). Specifically, we define a segment of ***x*** as consecutive measurements whose alignments to the *M* candidate squiggles (the columns of ***A***) are the same; see Fig. S5 for a toy example and Supplementary section 1.6 for our practical implementation of the segmentation. Suppose we have *N* segments in ***x***. Then, we compute the summary ***y*** = (*y*_1_,…, *y_N_*), where *y_i_* summarises information in ***x*** at its *i*-th segment. In this paper, we use medians for the summary as they are relatively robust to outliers.

We then compute the candidate squiggles and alignments of ***y*** (denoted by 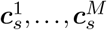 and ***A_s_***, respectively) from that of ***x***; see Supplementary section 1.6 for details. The NanoSplicer model for ***y*** uses 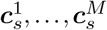 as model parameters and ***A***_*s*_ provides alignments between ***y*** and the model parameters.

#### 2.4.2 NanoSplicer model

For a given JWR, we build a mixture model to identify a splice junction among the *M* candidates. We introduce a latent variable *z* ∈ {1,…, *M*} indicating which candidate the junction squiggle came from. The mixture model for ***y*** = (*y*_1_,…, *y_N_*) can be written as

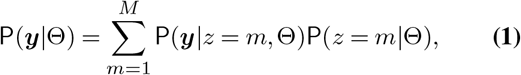

where 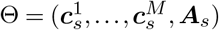. We assume that *y*_1_,…, *y_N_* are independent conditional on their means and standard deviations, yielding

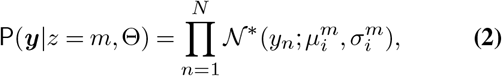

where 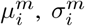 are the mean and standard deviation in the candidate squiggle 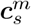 aligned to *y_n_* through ***A***_*s*_ (i.e., 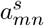). We model *y_n_* using modified normal distributions (denoted by 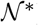) which have flat tails, making our method robust to measurements that match none of the *M* candidate squiggles; see Supplementary section 1.5 for details of these distributions. Such measurements could appear, for example, due to genetic variants which are not currently incorporated into our junction motifs (see section 2.2).

When there is other information reflecting the general propensity of a candidate to be a splice junction, we can model the mixing proportion 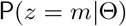 as a function of that information (e.g., nucleotide composition near splice sites for eukaryotes (Irimia and Roy, 2008); see section 4 and Supplementary section 1.8). Otherwise, 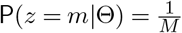.

#### 2.4.3 Identification of splice junctions

Identification of splice junctions can be performed by computing the posterior probability for each JWR:

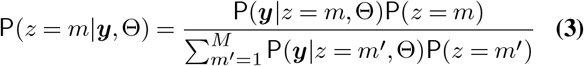

We call this the *assignment probability* that quantifies the support of the junction squiggle for each candidate. In practice, we restrict our identification to JWRs where a single candidate has strong support (e.g., we required an assignment probability > 0.8 in this paper).

#### 2.4.4 Squiggle information quality

In practice, we implement the following step to improve performance: the NanoSplicer model assumes that the junction squiggle corresponds to the location of the JWR, however Tombo can potentially align the JWR to an incorrect squiggle location. Therefore, we add a step to filter out junction squiggles that do not have a high quality alignment to any candidate squiggle, suggesting they emanate from an off-target read subsequence. First we measure the alignment quality between a junction squiggle and each of its candidate squiggles using the average log likelihood over the nucleotides of the candidate squiggle; see Supplementary section 1.7 for details. Then, we compute the maximum of these alignment qualities across *M* candidates, referred to as *squiggle information quality (SIQ)*. In practice, we restrict our splice junction identification to JWRs with SIQ bigger than a threshold to ensure their junction squiggles have high quality alignments at least one of their candidate squiggles. To choose a suitable threshold, we use an empirical distribution of SIQ constructed by pooling SIQ values from multiple JWRs in the analysis. Assuming that most JWRs are well aligned to correct squiggle locations, we choose an SIQ threshold that identifies junction squiggles whose SIQ values are much smaller than the majority of SIQs in the distribution. We illustrate our choice of thresholds in sections 3 and 4. See Supplementary section 2.9.1 for a discussion on how the choice of thresholds involves trade-offs between accuracy and the ability to identify splice junctions. Note that, although we propose this SIQ step to filter out junction squiggles from an off-target region, it can also help identify poor quality junction squiggles due to experimental artifacts (e.g., current spikes, pore blockages, or uneven dwell time of nucleotides in the pore, etc.) as they can also lead to poor alignments.

## 3 Synthetic RNA data analysis

A potential advantage of NanoSplicer is that it can exploit the information in squiggles. To assess the benefit of this feature, we compared the *accuracy*, defined as the proportion of correctly identified splice junctions, of NanoSplicer to the initial mapping results. We assessed the performance of NanoSplicer using *sequin* RNA standards (Hardwick *et al*., 2016), which enables us to compare NanoSplicer outputs to a known ground truth. Sequins are a set of synthetic spliced mRNA isoforms whose sequences and quantities are precisely known. An *in-silico sequin chromosome* contains each sequin gene and isoform, creating a known ground truth for the position of each splice junction, which mapping- and NanoSplicer-based results can be compared to. Sequins contain 160 isoforms from 76 genes and 745 splice junctions (all but 3 are canonical GT-AG junctions). We used a nanopore cDNA sequins dataset generated using the Oxford Nanopore Technologies GridION platform with a R9.4.1 MinION flowcell from Dong *et al*. (2021); see Supplementary section 2.1 for details of the data. We basecalled raw signals (squiggles) using Guppy 3.6.1 and mapped basecalled reads to the sequin genome using minimap2 (Li, 2018), resulting in 1,919,714 mapped reads to 76 genes and 4,270,674 JWRs. We deactivated the “splice flank” sequence preference option in minimap2 as this preference is not present in sequins. Supplementary sections 2.2 and 2.3 provides Guppy and minimap2 command lines for our analyses.

For the purpose of assessment, we first defined a ground truth splice junction for each JWR by choosing one among the known 745 sequin splice junctions as follows. We mapped the reads to the sequin isoforms using minimap2, providing a 1-1 correspondence between each read and a sequin isoform (Supplementary section 2.3.2). We restricted our assessment to 1,528,017 reads with the maximum mapping quality (mapQ = 60 in minimap2), for which we can accurately identify their corresponding isoforms. These reads contained 3,492,373 JWRs, of which 3,477,172 could be analysed by Nanosplicer (see Supplementary section 2.6 for details of the remaining 15,201 JWRs). Then, on the strand the isoform maps to, we searched for a sequin splice junction whose splice sites are within 10 bases of the mapped JWR splice sites and treated it as a ground truth for that JWR. All the JWRs have either one or zero known sequin splice junctions within 10 bases, supporting their correct assignment. The 134,529 JWRs (3.9%) without a nearby known splice junction have no ground truth and we will refer to them as *completely missed JWRs*.

### 3.1 NanoSplicer improves upon the initial mapping results using squiggles

We used 3,477,172 JWRs (encompassing 650 known sequin splice junctions) to assess the performance of NanoSplicer. The initial mapping failed to identify the ground truths for 228,181 JWRs, (6.56%, including 134,529 completely missed JWRs and 93,652 within 10 bases of the known sequin splice junction). Although any splice junctions identified by NanoSplicer for the completely missed JWRs will be incorrect, we include them in the analysis to assess how well SIQ (section 2.4.4) and assignment probability (section 2.4.3) in NanoSplicer recognise and filter out these JWRs.

We applied NanoSplicer to the dataset to improve upon the initial mapping results; see Supplementary section 2.12 for NanoSplicer run time. We chose candidates using the default option in section 2.2, used a uniform prior for mixing proportion (section 2.4.2), and chose −0.8 as a threshold for SIQ using an empirical distribution in Fig. S8. This meant all JWRs had the ground truth as a candidate, except for the completely missed JWRs, as well as JWRs where the ground truth was non-canonical and was not identified as the mapped splice junction. NanoSplicer reports identified splice junctions for 3,058,814 JWR, which have SIQ > −0.8 and strongest assignment probability > 0.8, with an accuracy of 96.4%. Therefore NanoSplicer improved the overall accuracy of splice junction detection compared to the initial mapping (93.4%).

### 3.2 NanoSplicer improvement is greatest when junction alignment quality (JAQ) is low

To better understand the advantages of NanoSplicer we next asked under what circumstances it improved upon splice junction detection in the initial mapping. Basecalling errors can result in low quality alignments between the JWR and the reference genome. Therefore we hypothesized that the initial mapping would perform poorly for JWRs with high basecalling errors and that the advantage of NanoSplicer will be greatest for these. We quantified the *junction alignment quality (JAQ)* for each JWR, which we defined as the percentage of matched bases in its alignment, to test this. For example, a JAQ of 0.96 can be interpreted as 4% of bases in an alignment being inserted/deleted or mismatched; see Supplementary section 2.5 for further details.

Fig. 2A shows the accuracy of each approach for JWRs with different ranges of junction alignment quality. NanoSplicer and minimap2 are both similarly accurate (98.3 vs 98.5%) when the JAQ is > 0.95 and the JWR sequence aligns almost perfectly. In such a circumstance there is little extra information to be obtained from the squiggle. However, at an alignment quality of 0.95 and below (51.2% of all JWRs), NanoSplicer improves upon the initial mapping accuracy, displaying progressively larger improvements as alignment quality decreases. For junction alignment qualities ≤ 0.8, NanoSplicer increased the raw accuracy by 21.0% and decreased the proportion of incorrect JWRs by 51 %. Furthermore, most multiexon genes have more than one splice junction and JAQ can vary over a read. We find that 17.6% and 76.8% of multi-exon reads have a JWR whose JAQ is ≤ 0.8 and 0.95 respectively. These results demonstrate that NanoSplicer has the potential to improve splice junction identification in a significant proportion of reads.

**Fig. 2.**
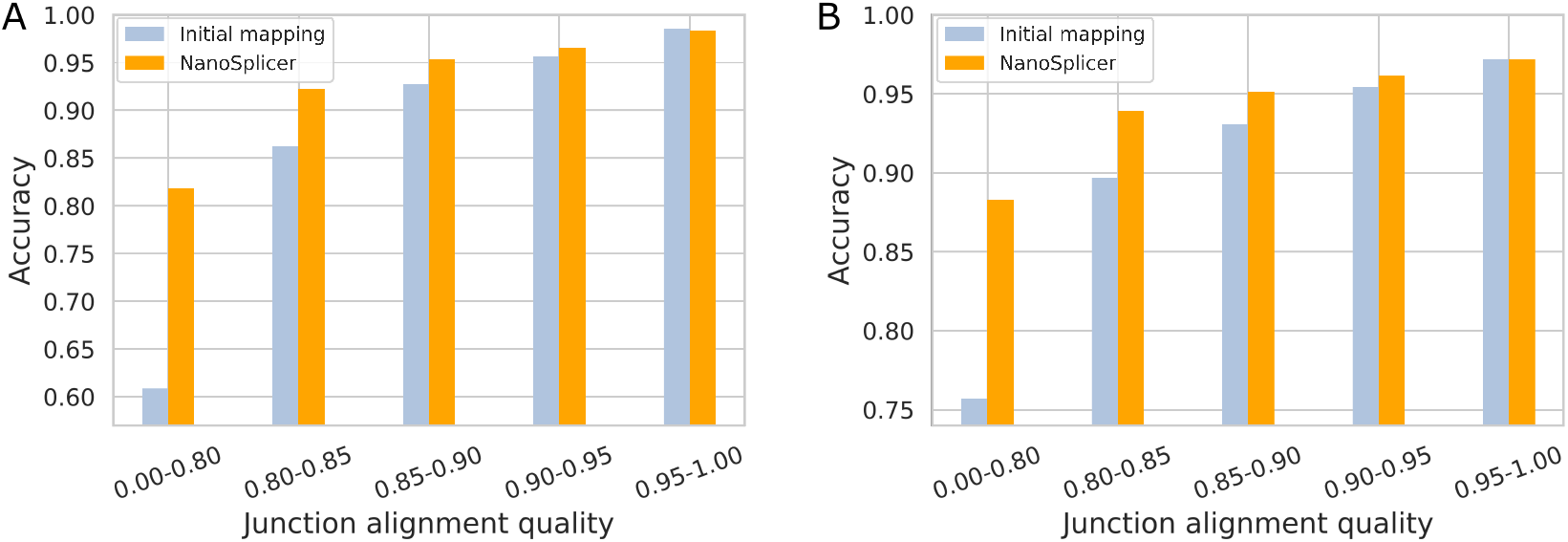
Splice junction identification accuracy of NanoSplicer and minimap2. A: Accuracy of splice junction identification in synthetic data. B: Accuracy of splice junction identification in biological data. A and B: all JWRs were binned based on junction alignment quality (JAQ). The interval “a-b” in the x-axis represents “a < JAQ ≤ b”. Initial mapping (from minimap2) is based on all JWRs. NanoSplicer accuracy is based on the JWRs where NanoSplicer identifies the splice junction (SIQ > −0.8, strongest assignment probability > 0.8). The number of JWRs in each JAQ bin, including completely missed JWRs, is shown in Table S1.

While the initial mapping reports splice junctions in all JWRs, NanoSplicer identified splice junctions for 3,058,814 JWRs and the accuracy of NanoSplicer is evaluated on this smaller set. Thus, multiple factors could contribute to its increased accuracy. These include the ability of SIQ and assignment probability to identify wrongly mapped JWRs, as well as NanoSplicer correction of the initial mapping results. To investigate the contributions of these factors, we compute the accuracy of the initial mapping on two sets of JWRs: 3,309,817 JWRs after SIQ filtering and 3,058,814 JWRs after SIQ and strongest assignment probability filtering (Supplementary section 2.7). All factors contribute to the increase in accuracy. NanoSplicer correction showed a clear benefit when JAQ ≤ 0.9. In contrast the contribution of SIQ filtering is proportionately large for low quality alignments (JAQ ≤ 0.8). Moreover, we observed that JWRs filtered by SIQ or assignment probability are enriched in completely missed JWRs (Supplementary section 2.8), supporting that these procedures help identify JWRs without true junctions as candidates.

### 3.3 Example of splice junction correction with NanoSplicer

To demonstrate how squiggles can provide extra information to identify splice junctions, Fig. 3 A1 & A2 shows an example JWR from the synthetic RNA dataset. Based on the ground truth of the synthetic RNA, the first 5 exonic bases after the 3’ splice site should be *“CCCAG”*, but were basecalled as *“TG”*. In the reference genome in Fig. 3A1, there are two “AG” 3’ splice motifs 5 bases apart, leading to two potential 3’ splice sites. The JWR was initially mapped to the wrong one 5 bases inside the exon due to the basecalling error. We compared the junction squiggle of this JWR to the candidate squiggles obtained from the true splice junction and the splice junction matching the initial mapping. Fig. 3A2 shows the alignments between the junction squiggle and the two candidate squiggles respectively. The squiggle from the true candidate splice junction is visually a closer match to the junction squiggle. NanoSplicer quantified this squiggle similarity, giving an assignment probability of 0.988 to the true candidate for this JWR. This example demonstrates how nanopore squiggle signal can be used to correct read alignments and accurately identify splice junctions. See Supplementary section 2.10 for more examples.

**Fig. 3.**
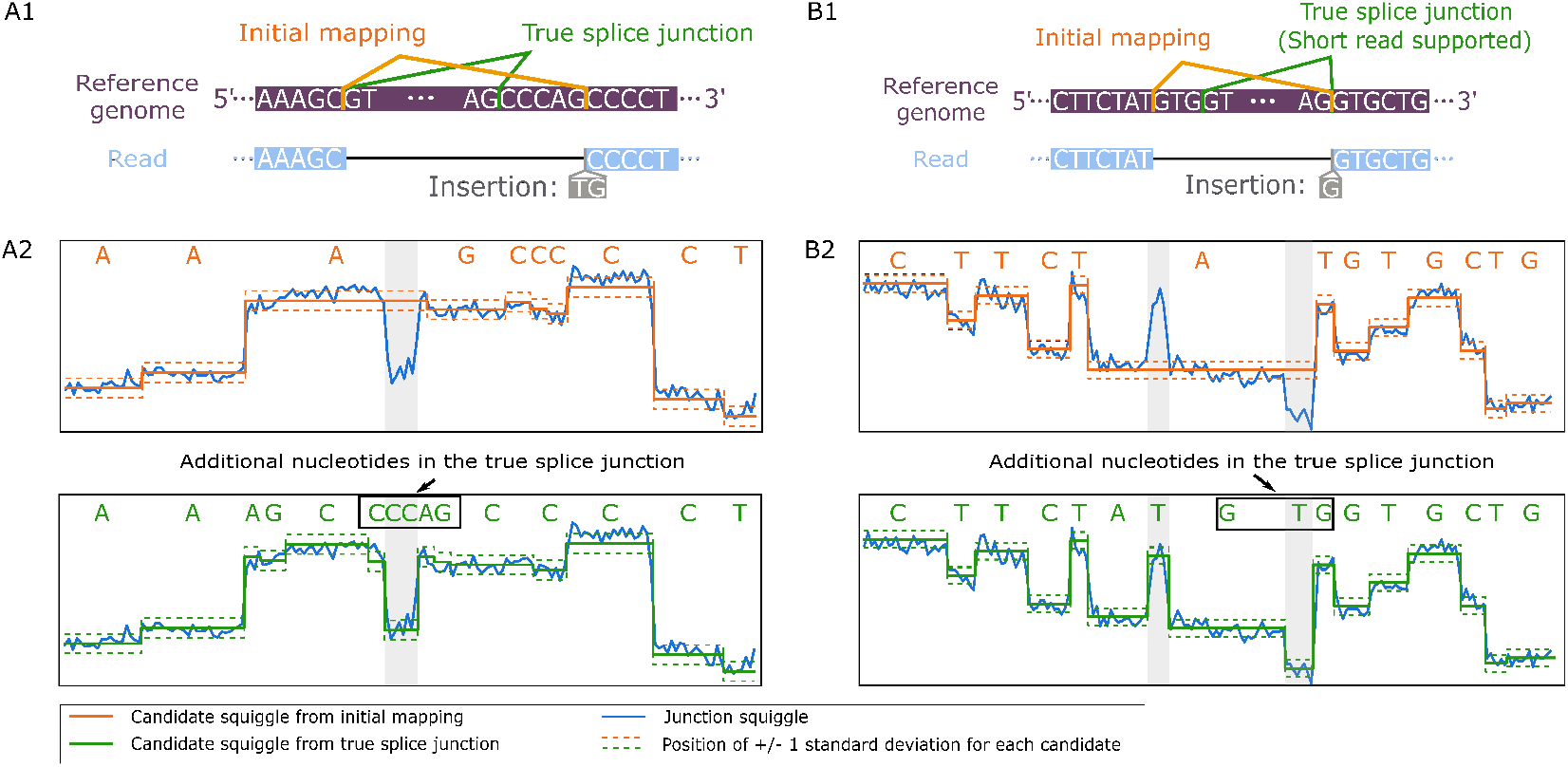
Examples of NanoSplicer correcting wrongly mapped JWRs. A1 & A2: Synthetic RNA data, B1 & B2: Biological data. A1 & B1: JWR mapping. Reference genome sequence shown in purple. Green line shows the location of the known ground truth splice junction. The mapped nanopore read (blue) shows the basecalled nucleotides
of the JWR and how they were aligned to the reference genome. Orange line shows the splice junction identified by the initial mapping of the JWR. Insertion - basecalled nucleotides in read that were not part of genome alignment. A2 & B2: Alignment between the junction squiggle for the JWR (blue) and the corresponding candidate squiggles from A1 & B1 (orange and green). Each junction squiggle current measurement is vertically aligned with its assigned mean-standard deviation in the candidate squiggle. The junction motifs for each candidate are shown at the top of each panel. Each nucleotide in the motifs is aligned with its corresponding squiggle position. Panels focus on the junction squiggle areas that distinguish between the candidates (grey background; the absolute difference in log likelihood between the two candidate models is bigger than 1.35; see Supplementary section 2.11 for details).

## 4 Biological RNA data analysis

In this section, we assess the performance of NanoSplicer on real biological data. To this end, we used a nanopore cDNA dataset (see Supplementary section 2.1 for detailed data description) generated from the lung cancer cell line NCI-H1975, for which short read data are also available (Holik *et al*., 2017). We basecalled all squiggles using Guppy 3.6.1 and mapped basecalled reads to the human GRCh38 assembly using minimap2. We focused our analysis on 758,330 reads mapped to chromosome 1, yielding 2,216,054 JWRs, of which 2,209,275 could be analysed by NanoSplicer (see Supplementary section 2.6 for details of the remaining 6,779 JWRs). Note that we retained the “splice flank” sequence preference option in minimap2 for this analysis, i.e., preferring “GTA” and “GTG” at the start of the intron and “CAG” and “TAG” at the end of the intron, as it improved mapping results.

For the purpose of assessment, we defined a ground truth for each JWR using the short read data from Holik *et al*. (2017), as short reads are expected to provide accurate information on the locations of splice junctions. First, we mapped the short reads to GRCh38 using STAR (Dobin *et al*., 2013) and considered splice junctions with at least 3 mapped short reads to be “known”; see Supplementary section 2.4 for details. Then, for each JWR, we searched for a known splice junction within 10 bases of the mapped splice sites and treated it as a ground truth for that JWR. To avoid ambiguity in determining the ground truth, we restricted our assessment to 2,073,181 JWRs that have at most one nearby known splice junction.

### 4.1 NanoSplicer improves upon the initial mapping results using squiggles

We assessed the performance of NanoSplicer on the 2,073,181 JWRs; see Supplementary section 2.12 for the run time. As previously, we compared the accuracy (i.e., the proportion of correctly identified splice junctions) of NanoSplicer to the initial mapping results. The initial mapping failed to identify the ground truths for 111,560 JWRs (5.4%), including 73,043 completely missed JWRs. In this analysis, we incorporated “splice flank” sequence preferences, like those in minimap2, as prior mixing proportions in NanoSplicer; see Supplementary section 1.8 for details. Other steps in the NanoSplicer workflow were similar to the synthetic data analysis: candidates were chosen using the default option in section 2.2, and SIQ > −0.8 (see Fig. S8) and strongest assignment probability > 0.8 thresholds were used. NanoSplicer identified splice junctions for 1,902,248 JWRs with an accuracy of 96.1%, confirming it improves splice junction detection from biological data.

Similar to the synthetic data analysis, the improvement in splice junction identification with NanoSplicer increased as junction alignment quality (JAQ) decreased (Fig. 2B). NanoSplicer improved upon the initial mapping when JAQ ≤ 0.95 and was particularly pronounced below a JAQ of 0.8. For a JAQ ≤ 0.8, NanoSplicer increases the accuracy by 12.6% and halved the proportion of incorrect JWRs. As the accuracy of NanoSplicer is evaluated on a smaller set of JWRs (1,902,248) than the initial mapping, we again investigated the contributions of SIQ, strongest assignment probability and NanoSplicer correction to the increased accuracy. Supplementary section 2.7 revealed results consistent with the synthetic data analysis: all factors contribute to the increased accuracy; NanoSplicer correction is beneficial when JAQ ≤ 0.95 and SIQ filtering has a relatively larger contribution when JAQ ≤ 0.8.

### 4.2 Example of splice junction correction with NanoSplicer

Fig. 3 B1 & B2 shows an example JWR from the biological dataset. In Fig. 3B1 there are two “GT” 5’ splice motifs only 3-bases apart at the 5’ splice site, leading to two candidate splice junctions. One of the candidate splice junctions is supported by the short read data and is assumed to be the true one. However, the nanopore read in Fig. 3B1 was mapped to the other candidate splice junction due to basecalling errors at the 5’ splice site (the *“GTG”* bases preceding the true splice junction were basecalled as only *“G”* in the nanopore read). We compared the junction squiggle of this JWR to the squiggles obtained from both candidate splice junctions. Fig. 3B2 shows the alignments between the junction squiggle and the two candidate squiggles respectively. The shape of the squiggle for the true candidate is clearly a better match for the junction squiggle, while the initial mapping candidate misses clear signal changes indicative of additional bases. NanoSplicer quantified this squiggle similarity, leading to an assignment probability of 0.997 to the true candidate for this JWR. See Supplementary section 2.10 for more examples.

### 4.3 Identification of non-canonical junctions

A small proportion of splice junctions do not use the canonical GT-AG motif and are challenging to identify from error prone reads. Comparing the JWRs from the NCI-H1975 long-reads to the short-read data revealed 10488 (0.47%) had a non-canonical truth. Minimap2 correctly identified 3993 of these (38.1%), while NanoSplicer identified 3572 of 8626 (41.4%) after SIQ and assignment probability filtering. Using the default option NanoSplicer can only consider non-canonical junctions as candidates if they are the mapped splice junctions, meaning it cannot correct the 6495 JWRs where minimap2 was incorrect. NanoSplicer also allows user supplied lists of candidate splice junctions as inputs. We added human non-canonical splice junctions from RefSeq (O’Leary *et al*., 2016) and re-performed the NanoSplicer analysis. NanoSplicer now correctly identified 9344 of 9625 (97.1%) of the JWRs with a non-canonical truth that passed filtering. These results demonstrate the flexible inputs Nanosplicer can utilise and how Nanosplicer performance for non-canonical splice junction identification can benefit from their use.

## 5 Conclusion

We have developed a novel method, NanoSplicer, to accurately identify splice junctions using nanopore sequencing. The method, adapting the “squiggle matching” idea, exploits the information in squiggles to improve identification. This enables NanoSplicer to identify splice junctions solely from the nanopore data without requiring annotations or matched short reads. It also enables its performance to be independent of other reads or read depth, having the potential to better identify rare splice junctions (Supplementary section 2.13). Using both synthetic and real data, we show that NanoSplicer improves upon the initial mapping, particularly when the basecalling error rate near splice junctions is high, demonstrating the contribution of squiggle information to splice junction identification.

To our knowledge, this is the first method that exploits squiggle information for splice junction identification. Therefore, there are many opportunities for potential improvements. First, the NanoSplicer model treats the summary values of junction squiggles as observed data and ignores their uncertainty, possibly leading to a decreased accuracy. One potential way to incorporate the uncertainty is to exploit the likelihood approximation as described in Shim *et al*. (2021), yielding likelihoods expressed by the estimates of model parameters and their standard errors. Second, very short or long dwell times (i.e., the duration of a translocation event) may not reflect typical translocation events (Díaz Carral *et al*., 2021), potentially causing misleading results. Here, we partly address this issue by filtering out summary values based on a very small or large number of measurements, but a more principled approach, such as modelling dwell time, could potentially improve performance. Third, here we predict expected squiggles from junction motifs using the “expected current level model” in Tombo (Stoiber *et al*., 2016), but this can be achieved by using other models that are appropriate for the chemistry in use. Indeed, the optimal choice of models may be one learned from the data at hand. Finally, NanoSplicer has been comprehensively tested only for analysis of Nanopore cDNA data and the R9 pore but will be tested on direct RNA sequencing and the R10 pore in the future.

NanoSplicer identifies splice junctions only among candidates, potentially leading to false detection when the true junctions are not included. We are not alone in having this limitation; for example, other tools restrict their correction to junctions from annotations or matched short reads (Tang *et al*., 2020; Wyman and Mortazavi, 2019), and/or to junctions supported by mapped reads (Kovaka *et al*., 2019; Kuo *et al*., 2020; Parker *et al*., 2021). However, NanoSplicer provides flexible options for candidate selection (sections 2.2 and 4.3), enabling users to use context-dependent candidates. Moreover, our empirical analysis in Supplementary section 2.8 shows that SIQ and assignment probability help filter out JWRs without true junctions as candidates, reducing false identifications.

NanoSplicer has been designed and tested for accurate identification of splice junctions. The identified junctions could be leveraged in different types of downstream analyses. For some analyses, however, NanoSplicer’s outputs should be used with care as it identifies splice junctions only for JWRs whose squiggles are informative. For example, if the outputs are used for splice junction quantification, excluding JWRs without identified outputs or correcting them using splice junctions from other reads may lead to less accurate quantification because they are not a random subset of all JWRs (Fig. S15). Additionally, utilising mapped junctions from these JWRs to supplement NanoSplicer outputs would decrease overall accuracy as such JWRs tend to have lower JAQs (Fig. S16). We are currently investigating to what extent these ad hoc approaches can provide good performance in splice junction or isoform quantification.

Our analysis shows that NanoSplicer improvement is greatest when junction alignment quality (JAQ) is low, while initial mapping results tend to be correct for JWRs with high JAQs because their sequences align almost perfectly. In practice, the first step in our software is calculation of JAQs for JWRs, which provides useful and quickly accessible information on long-read junction quality for an experiment or read of interest. Thus, our software offers an option to output JAQs with-outrunning the identification step. Additionally, NanoSplicer provides an option to run it on JWRs below a user-specified JAQ threshold (default 0.95) to reduce its run time and focus on the JWRs it is most likely to correct. Nanopore sequencing accuracy is increasing over time, however even as median read accuracy has increased, Nanopore read accuracy distributions still exhibit a long tail of reads with lower accuracy. Therefore, there remains a significant proportion of reads for which splice junction identification (and subsequent isoform identification) can be enabled by NanoSplicer.

## Acknowledgements

We thank Luyi Tian and Quentin Gouil for generating the lung cancer data and Matthew Ritchie for allowing the use of this data; Terry Speed, Martin Smith, and James Ferguson for invaluable discussion; and Yao-ban Chan for helpful comments on a draft manuscript. We also thank the members of the Shim, Clark, and Speed labs and the funwip group for helpful discussions.

## Supplementary material

Supplementary material is available at https://github.com/shimlab/NanoSplicer/blob/master/docs/NanoSplicer_Supplementary.pdf.

